# *Paenibacillus xylinteritus* sp. nov., a novel bacterial species isolated from grapevine wood

**DOI:** 10.1101/2022.12.09.519748

**Authors:** Rana Haidar, Livio Antonielli, Stéphane Compant, Ursula Sauer, Caroline Pandin, Claire Gassie, Amira Yacoub, Maria Chrysovergi, Eléonore Attard, Patrice Rey, Rémy Guyoneaud

## Abstract

Bacteria naturally colonize grapevine wood, and some populations interact synergistically with fungal pathogens to promote wood degradation. In this study, we characterized a bacterial strain, designated S150^T^, isolated from the wood of grapevine cultivar Sauvignon Blanc. The strain was previously known as promoting wood degradation caused by the causal agent of white-rot, *Fomitiporia mediterranea*. Phylogenetic analysis based on the full-length 16S rRNA gene sequence showed that the strain S150^T^ belonged to the genus *Paenibacillus*. Based on 16S rRNA gene sequences, S150^T^ shared low sequence similarity values (between 96.5 and 97.5%) with closed recognized members of the genus (*P. jilunlii* Be17, *P. tepidiphilus* SYSU G01001, *P. tengchongensis* SYSU-G-01003, *P. sonchi* X19-5, *P. helianthi* P26E). The cell wall peptidoglycan of S150 contains meso-diaminopimelate. MK6 and MK7 are the two respiratory quinones. The main cellular fatty acids are iso-C_16:0_ and anteiso-C_15:0_. The polar lipids are diphosphatidylglycerol, phosphatidylethanolamine, three amino phospholipids, two phospholipids and three polar lipids. The whole-genome is 7.45 Mb, with a G+C content of 52.54%. The average nucleotide identity (ANI) value between S150^T^ and the closely related *Paenibacillus* member (*P. sonchi* X19-5) was 85.06%. Biochemical and physiological analyses revealed that strain S150^T^ has different characteristics from reference strains *Paenibacillus* spp.. Phenotypic, phylogenetic and chemotaxonomic analyses show strain S150^T^ is a novel species of the genus *Paenibacillus*, for which the name *Paenibacillus xylinteritus* is proposed.

---

Grapevines are naturally colonized by diverse communities of epiphytic and endophytic microorganisms in various organs such as root, stem, wood, flowers, leaves and berries. These microorganisms impact grape and wine production [1–3]. Theses bacteria could have beneficial, neutral or pathogenic effect on grapevine [4–7]. Recently Haidar *et al*. [7] isolated 237 grapevine-wood bacterial strains from decaying plants of Sauvignon Blanc cultivar. One strain that interacted synergistically with the fungal wood-pathogen *Fomitiporia mediterranea* was selected for its ability to promote grapevine wood degradation [7]. This strain, isolated from cordon wood tissues and designated as S150^T^, corresponded to a member of the genus *Paenibacillus*.

Many species of the genus of *Paenibacillus* were reported to colonize different organs of grapevine [8–11]. While some *Paenibacillus* species display biocontrol activities against fungal pathogens of grapevine such as *Botrytis cinerea, Phaeomoniella chlamydospora* and *Neufosicoccum parvum* [4–5]. Other species develop synergistic relationships with *B. cinerea* and/or *N. parvum* that favor their pathogenic process [4]. *In natura, Paenibacillus* species have been isolated from various environments, including soil, water, insect, rhizosphere, food, plants and clinical samples [12]. Members of this genus are rod-shaped, sporulating and facultative anaerobes and *Paenibacillus* genus contains more than 260 recognized species (http://www.bacterio.net/paenibacillus.html).

S150^T^ was isolated from wood tissue of the cordon of grapevines (cultivar Sauvignon Blanc, *Vitis vinifera* L.) grafted onto the rootstock 101-14 MGt. The used medium was R2A agar medium amended with 100 mg.L^-1^ cycloheximide. The experimental vineyard was located near Bordeaux (INRAE, Villenave d’Ornon, France, 44°47’24.8”N, 0°34’35.1”W) [7].

Genomic DNA was extracted from pellets obtained after the centrifugation of pure cultures grown for 48h in Tryptone-Soy Broth (TSB, Biokar) using the cetyltrimethylammonium bromide (CTAB) chloroform/isoamyl alcohol (24:1) protocol. The 16S rRNA gene was amplified and the obtained sequence was merged manually based on alignment from MUSCLE [13]. Phylogenetic analysis, comprising a number of different strains of *Paenibacillus* species, and based on 16S rRNA gene sequences, was constructed.

Phylogenetic analysis based on the full-length 16S rRNA gene sequence and sequence comparisons with the 16S rRNA gene sequences stored in GenBank and the GTDB-Tk databases revealed that the strain S150^T^ has high sequence similarity with the members of the genus *Paenibacillus*. Based on 16S rRNA gene sequence and phylogenetic analysis, *P. jilunlii* Be17, *P. tepidiphilus* SYSU G01001, *P. tengchongensis* SYSU-G-01003 [17], *P. helianthi* P26E [18], *P. piscarius* P121 [19], *P. riograndensis* SBR5 [20], *P. sonchi* X19-5 [15], *P. graminis* RSA19 [21], *P. salinicaeni* LAM0A28 [22], *P. azotifigens* NF2-4-5 [23] were selected as the closest recognized neighbors of strain S150^T^.

The position of the novel strain S150^T^ in relation with other members of the genus *Paenibacillus* was shown in the phylogenetic tree (Figure 1). It indicated that the bacterial strain investigated in this study formed a distinct monophyletic clade from the genus *Paenibacillus*. Strain S150^T^ shared low sequence similarity with the type strains of: *P. jilunlii* Be17 GQ985393^T^ (97.5%) [14], *P. sonchi* X19-5^T^ DQ358736 (97.2%) [15], *P. tepidiphilus* SYSU G01001^T^ (96.7%) [16], suggesting that strain S150^T^ should be classified as a novel species in the genus *Paenibacillus*.

**Figure 1.**
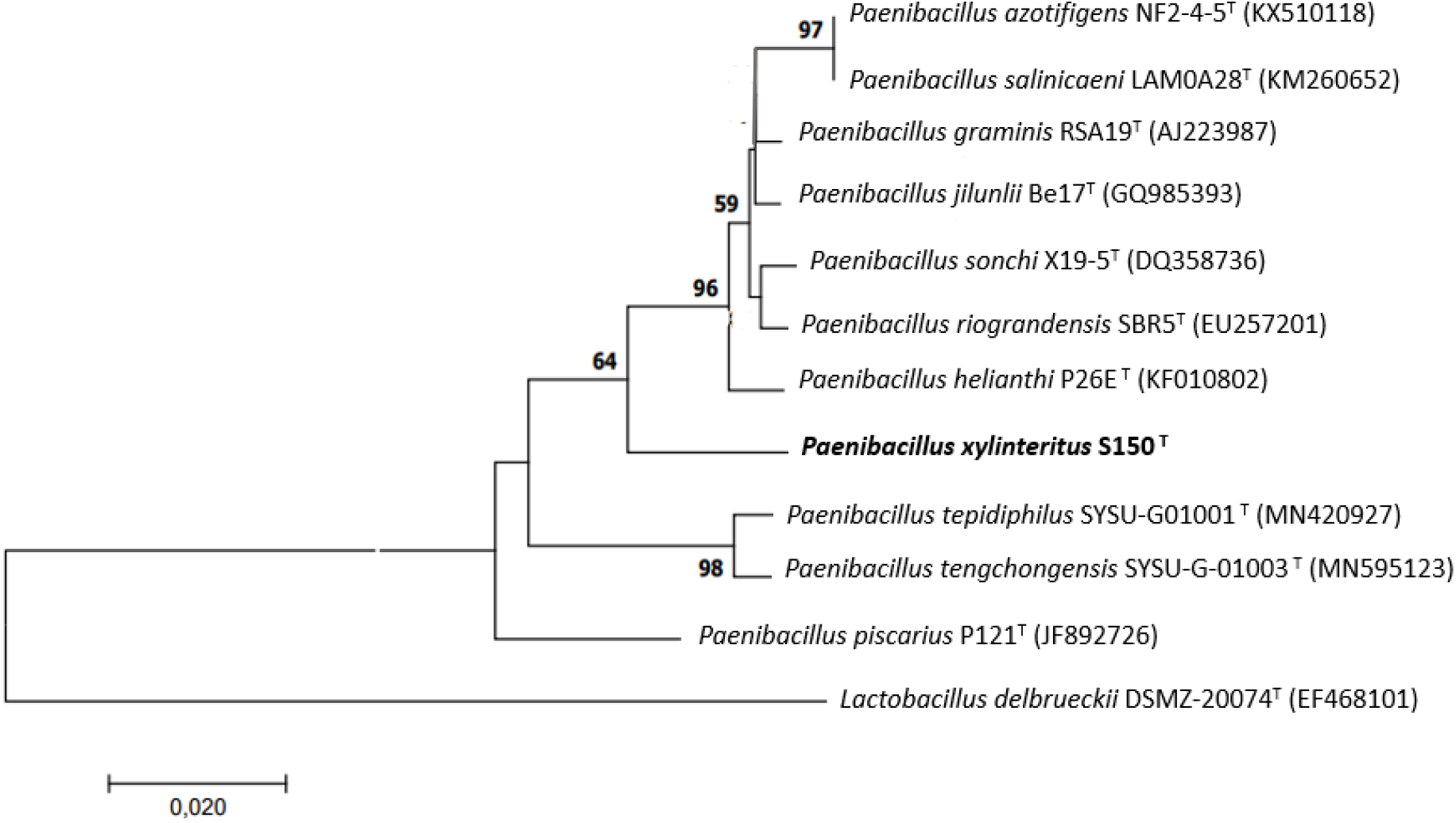
Maximum-likelihood phylogenetic tree based on 1342 bp showing the phylogenetic relationships of strain S150^T^ and the closely related members of the genus *Paenibacillus*. Bootstrap values (expressed as percentages of 1000 replications) greater than 50% are shown at branch points. *Lactobacillus delbrueckii* DSMZ 20074^T^ (EF468101) was used as an outgroup. Scale bar, 0.02 substitutions per nucleotide position.

The whole-genome shotgun sequencing of S150^T^ was performed on an Illumina NovaSeq 6000 mode S2 (GATC Biotech, Konstanz, Germany), producing approximately 6.2 million paired-end reads of 150 bp length. After quality filtering and adapter stripping with fastp v0.20.1 [24], 5.8 M (94%) high quality sequences remained, accounting for 1.7 G bases. After filtering, 93.6% of remaining reads showed an average Phred quality score of Q30. Filtered reads were then assembled in contigs with SPAdes v3.14.0 [25]. Contigs shorter than 500 bp and below 2X coverage were discarded. Reads were mapped against contigs using Bowtie 2 v2.3.4.3 [26] and the BAM file was inspected with QualiMap v2.2 [27], showing a mean coverage of 229.36X, a GC content of 52.54% and a mean mapping quality of 41.12. Contigs were aligned against the entire NCBI nt database with BLAST v2.10.0 [28] and the hit table was processed using BlobTools v1.1.1 [29]. After inspecting the BLAST results, contigs not assigned to *Paenibacillus* spp., were not considered in the follow-up analysis. Genome assembly evaluation was performed with QUAST v5.0.0 [30], resulting in 102 high-quality contigs, with a total size of 7.42 Mb and a final GC content of 52.35%. The largest contig was 649,111 bp and the N50 was 144,333. Genome quality assessment displayed a completeness of 99.2% while using BUSCO v4.0.6 [31] with 124 marker genes (Bacteria odb10). While using 468 single-copy orthologous genes with CheckM v1.0.18 [32], a completeness of 99.8% was reported and no significant evidence of contaminant contigs. Phylogenetic placement analysis with GTDB-Tk v1.7.0 [33] (database release 202) confirmed that the sequenced strain belongs to *Paenibacillu*s genus, with no further identification at species level.

The ncbi-genome-download v0.3.1 resource was used to download the genome sequence of 823 available *Paenibacillus* spp. from GenBank repository (April 2022). Downloaded FASTA files served as input for the average nucleotide identity (ANI) analysis accomplished with the BLAST approach, using pyani [34] v0.2.11. The closest match to *Paenibacillus* sp. S150 genome sequence was *P. sonchi* X19-5T ^T^, with a similarity percentage of 85.06%.

Gene prediction and annotation, performed with PROKKA [35] v.1.14.5, identified 6,292 genes among which 6,200 were coding sequences (CDS). A total of 6 rRNA and 85 tRNA were predicted. In-silico antimicrobial resistance screening was carried out with ABRicate v1.0.0, showing the presence of an rphC gene (rifamycin-inactivating phosphotransferase). The sequence hit showed an identity of 82.45% and coverage 99.85% against the NCBI AMR [36] database, confirmed by analogous search in MEGARES database [37]. No virulence genes were detected. Plasmid search was performed using Platon v1.16 [38], showing the presence of no additional replicons. 230 genes involved in carbohydrate transport and metabolism were detected using the ClassicRAST [39]. In addition, seven genes involved in xylose (xyloside) degradation were detected confirming the potential role of this strain in grapevine wood degradation. The genome has genes encoding proteins predicted to be involved in the plant-microbe interaction, including motility and chemotaxis, siderophore, biofilm, and iron as well as genes related to oxidative stress and detoxification.

Colony morphology and cell morphology were determined by light microscopy on cultures grown on TSA. A simple and rapid (KOH) method was used for the determination of the Gram type. Strain S150^T^ was visualized by atomic force microscopy (AFM, Figure 2). Pure cultures were centrifuged at 4,500 rpm for 10 minutes at a temperature of 4°C, then the medium was removed, bacteria were washed in distilled water twice and finally resuspended in 5 mL distilled water. Ten μL were placed on glass slides and dried under ambient, sterile conditions for about 0.5 hours before conducting AFM scans.

**Figure 2.**
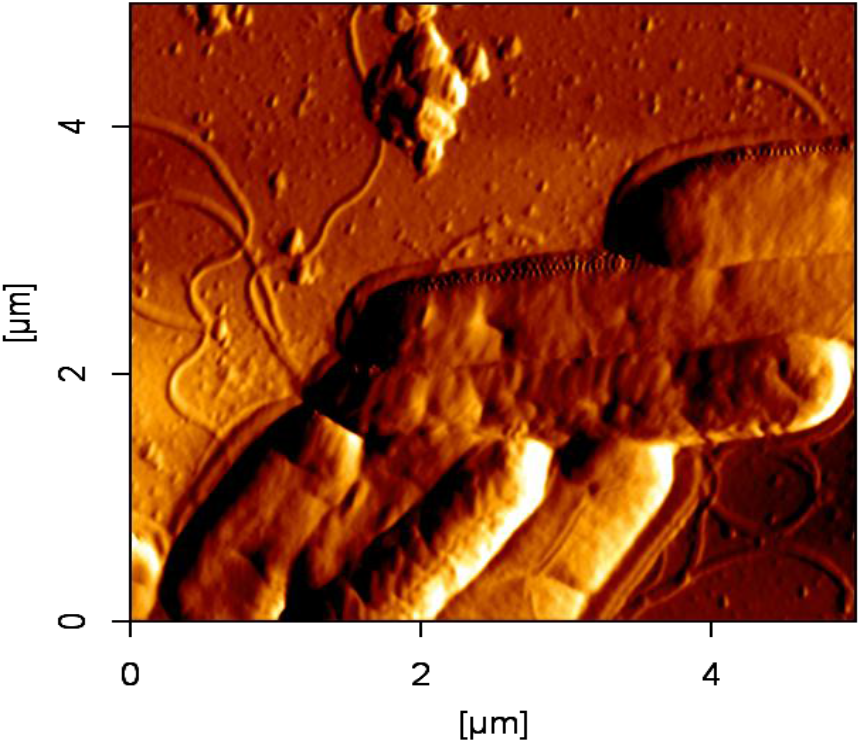
AFM Micrographs of cells on glass with flagella and the typical smooth appearance of a Gram-positive strain.

Scans were performed with AFM Nanowizard II (JPK Instruments, Berlin, Germany) in Intermittent Contact Mode in air using HQ-NSC35 /AlBS tip from μMasch (Wetzlar, Germany) at 300 kHz drive frequency. For high resolution scans the z-range was set to 3 μm, the line rate was typically 2.3 - 0.4 Hz. Data processing was performed with JPK data processing software 5.0.133. The catalase activity was determined by the production of bubbles when bacterial cells were placed in contact with 3 % (v/v) H_2_O_2_ solution. The oxidase activity was evaluated with dimethyl-p-phenylenediamine discs. The effect of temperature (4, 20, 30, 37and 42 °C) on cell growth of strain S150^T^ and its growth in the presence of NaCl (0-5 %) were determined in TSB medium. The substrate utilization and the enzyme production tests were performed using API 20 (API; bioMerieux) and GEN III Micro Plate (Biolog) assays according to the manufacturer’s instructions.

Cells of strain S150^T^ were Gram positive, slightly motile and rod shaped with flagella (Figure 2). Colonies of strain S150^T^ grown on TSA for 48h were small, variable, opaque and creamy. S150^T^ tolerated NaCl up to 3% (w/v) (growth optimal in the absence of NaCl) and grew at 20-37°C (optimum 30 °C).

Using Biolog GEN III, strain S150^T^ was not able to reduce nitrate to nitrate, to produce urease, indole and arginine dehydrolase. But this strain produced β-glucosidase, protease, gelatinase, β-galactosidase and used D-glucose, D-mannitol, N-acetyl-D-glucosamine, D-maltose and potassium gluconate.

Analysis of the cellular fatty acid, polar lipids, respiratory quinone and peptidoglycan were carried out by DSMZ services, Leibniz-institut DSMZ-Deutsche Sammlung von Mikroorganismen und Zellkulturen GmbH, Braunschweig, Germany. The Biomass for fatty acid analysis was harvested from cultures grown on TSA for 2days at 28C°. The isolation of the cell wall peptidoglycan and the study of its structures were carried out according to the protocols described by Schumann [40]. The cell wall peptidoglycan of S150 was A1γ’meso-Dpm-direct. Extraction and HPLC-DAD analysis showed the presence of menaquiones (MK): MK6 (3.1 %) and MK7 (96.9%).

The polar lipids of S150 were diphosphatidylglycerol (DPG), phosphatidylethanolamine (PE), three amino phospholipids (APL), two phospholipid (PL) and three polar lipid (L). The major one of strain S150 (>5%) was anteiso-C_15:0_ (47.3 %). Other fatty acides detected in S150 were iso-C_16:0_ (19.0 %), C_16:0_ (12.5 %) and iso-C_14:0_ (8.6%) which was consistent with the major fatty acids profile of other species of the genus *Paenibacillus* [23].

Detailed differentiating features of S150^T^ and the closest type strains of selected *Paenibacillus* species are listed in Table 1.

**Table 1:**
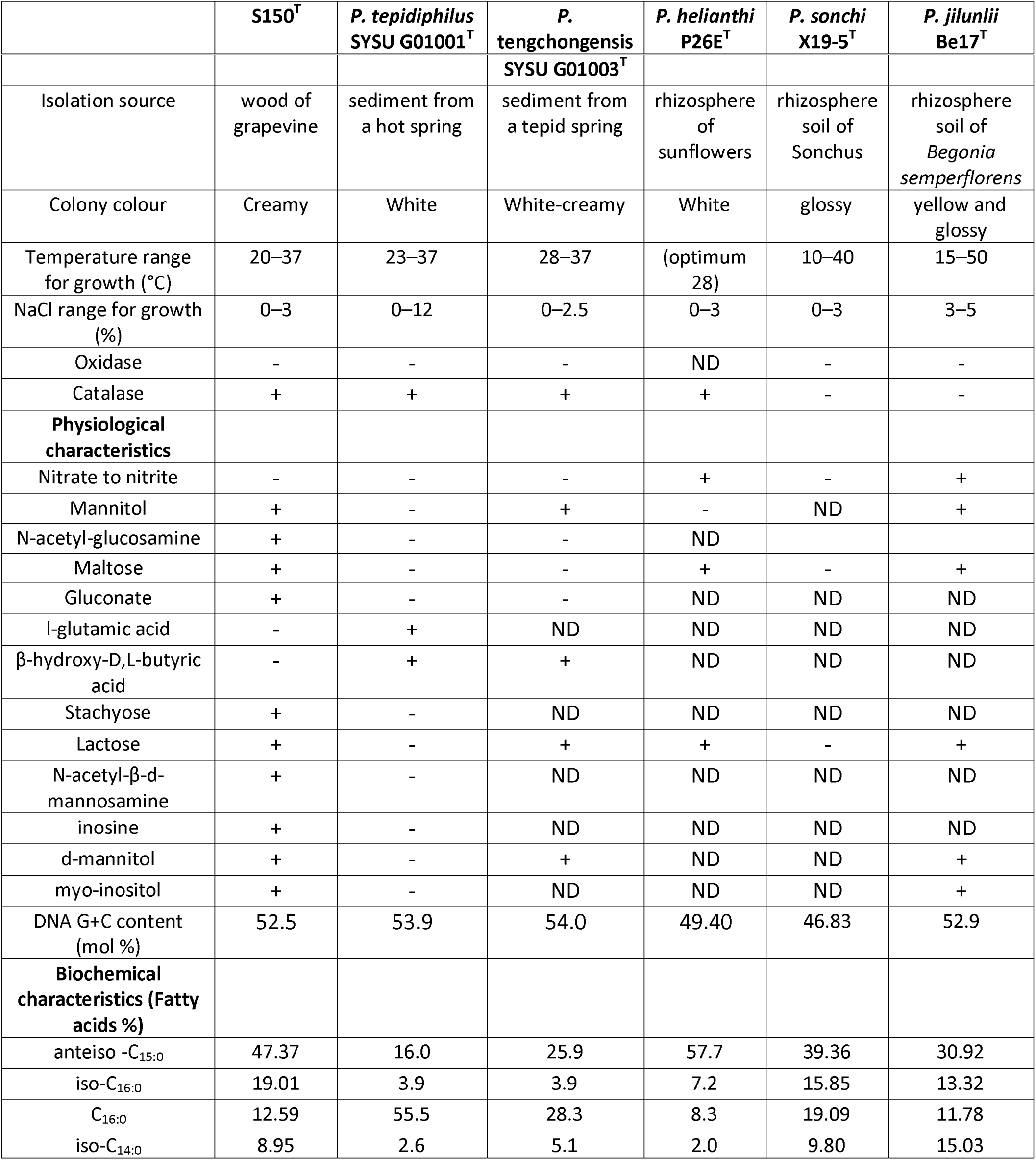
Different characteristic features among S150 and the closely related type strains of strains of selected *Paenibacillus* species.

|  |  |  |  |  |  |  |
| --- | --- | --- | --- | --- | --- | --- |
|  | **S150<sup>T</sup>** | ***P. tepidiphilus***<br>**SYSU G01001<sup>T</sup>** | ***P.***<br>**tengchongensis** | ***P. helianthi***<br>**P26E<sup>T</sup>** | ***P. sonchi***<br>**X19-5<sup>T</sup>** | ***P. jilunlii***<br>**Be17<sup>T</sup>** |

|  |  |  | **SYSU G01003<sup>T</sup>** |  |  |  |
| --- | --- | --- | --- | --- | --- | --- |
| Isolation source | wood of grapevine | sediment from a hot spring | sediment from a tepid spring | rhizosphere of sunflowers | rhizosphere soil of *Sonchus* | rhizosphere soil of *Begonia semperflorens* |
| Colony colour | Creamy | White | White-creamy | White | glossy | yellow and glossy |
| Temperature range for growth (°C) | 20–37 | 23–37 | 28–37 | (optimum 28) | 10–40 | 15–50 |
| NaCl range for growth (%) | 0–3 | 0–12 | 0–2.5 | 0–3 | 0–3 | 3–5 |
| Oxidase | - | - | - | ND | - | - |
| Catalase | + | + | + | + | - | - |
| **Physiological characteristics** |  |  |  |  |  |  |
| Nitrate to nitrite | - | - | - | + | - | + |
| Mannitol | + | - | + | - | ND | + |
| N-acetyl-glucosamine | + | - | - | ND |  |  |
| Maltose | + | - | - | + | - | + |
| Gluconate | + | - | - | ND | ND | ND |
| l-glutamic acid | - | + | ND | ND | ND | ND |
| β-hydroxy-D,L-butyric acid | - | + | + | ND | ND | ND |
| Stachyose | + | - | ND | ND | ND | ND |
| Lactose | + | - | + | + | - | + |
| N-acetyl-β-d-mannosamine | + | - | ND | ND | ND | ND |
| inosine | + | - | ND | ND | ND | ND |
| d-mannitol | + | - | + | ND | ND | + |
| myo-inositol | + | - | ND | ND | ND | + |
| DNA G+C content (mol %) | 52.5 | 53.9 | 54.0 | 49.40 | 46.83 | 52.9 |
| **Biochemical characteristics (Fatty acids %)** |  |  |  |  |  |  |
| anteiso -C<sub>15:0</sub> | 47.37 | 16.0 | 25.9 | 57.7 | 39.36 | 30.92 |
| iso-C<sub>16:0</sub> | 19.01 | 3.9 | 3.9 | 7.2 | 15.85 | 13.32 |
| C<sub>16:0</sub> | 12.59 | 55.5 | 28.3 | 8.3 | 19.09 | 11.78 |
| iso-C<sub>14:0</sub> | 8.95 | 2.6 | 5.1 | 2.0 | 9.80 | 15.03 |

In summary, the above phenotypic, phylogenetic and chemotaxonomic results showed that our strain S150 represents a novel species of the genus *Paenibacillus*, for which the name *Paenibacillus xylinteritus* sp. nov. is proposed.

## DESCRIPTION OF *PAENIBACILLUS XYLINTERITUS* SP. NOV

*Paenibacillus xylinteritus* (xyl’in.te.ri.tus Gr. prefix *xylo* wood; L. masc. n. *interitus* destruction; N.L. masc. n. *xylinteritus* the destruction of wood)

Cells are Gram positive, rod-shaped, slightly motile with flagella. Colonies grown TSA agar medium are variable, opaque and creamy. The temperature range for growth is 20–37 with optimal growth at 30 °C. The bacteria grow in the presence 3 % (w/v) NaCl but not at 3.5%. Catalase- and oxidase-negative. In API and 20NE, negative for oxidase, urease, arginine dihydrolase, lysine décarboxylase, ornithine décarboxylase, indole production and nitrate reduction but positive for β-glucosidase, protease, gélatinase, β-galactosidase, D-glucose, D-mannitol, N-acetyl-D-glucosamine,D-maltose and potassium gluconate.

In Biolog GEN III MicroPlate is positive for D-maltose, D-trehalose, D-cellobiose, gentiobiose, sucrose, D-turanose, stachyose, D-raffinose, α-D-lactose, D-melibiose, β-methyl-D-glucoside, D-salicin, N-acetyl-D-glucosamine, N-acetyl-β-D-mannosamine, α-D-glucose, D-mannose, D-fructose, D-galactose, L-fucose, L-rhamnose, inosine, D-mannitol, myo-inositol, glycerol, pectin, D-galacturonic acid, L-galactonic acid lactone, D-gluconic acid, glucuronamide, acetoacetic acid and aetic acid. The strain doesn’t use N-acetyl-D-galactosamine, N-acetyl neuraminic acid, D-arabitol, D-glucose-6-PO4, D-aspartic acid, D-serine, gelatin, glycyl-L-proline, L-alanine, L-arginine, L-aspartic acid, L-glutamic acid, L-histidine, L-pyroglutamic acid, L-serine, D-glucuronic acid, mucic acid, quinic acid, D-saccharic acid, p-hydroxy-phenylacetic acid, methyl pyruvate, D-lactic acid methyl ester, L-lactic acid, citric acid, α-keto-glutaric acid, D-malic acid, L-malic acid, bromo-succinic acid, tween 40, γ-amino-butryric acid, α-hydroxy-butyric acid, β-hydroxy-D,L-butyric acid, α-keto-butyric acid, propionic acid, formic acid, maltose, trehalose, gentiobiose, sucrose, turanose, melibiose, glycerol, l-glutamic acid, pectin, d-gluconic acid, methyl pyruvate, β-hydroxy-d,l-butyric acid and acetic acid but negative for stachyose, lactose, N-acetyl-β-d-mannosamine, N-acetyl-d-galactosamine, N-acetyl neuraminic acid, inosine, d-sorbitol, d-mannitol, d-arabitol, myo-inositol, d-aspartic acid, d-serine, glycyl-l-proline, l-arginine, l-aspartic acid, l-histidine, l-pyroglutamic acid, l-serine, mucic acid, quinic acid, d-saccharic acid, p-hydroxyphenylacetic acid, d-lactic acid methyl ester, citric acid, α-ketoglutaric acid, d-malic acid, l-malic acid, bromosuccinic acid, γ-aminobutryric acid, α-hydroxybutyric acid, α-ketobutyric acid, acetoacetic acid, propionic acid and formic acid. The cell wall contains meso-diaminopimelate (A1γ’meso-Dpm-direct). MK6 and MK7 were the two respiratory quinones. The main cellular fatty acids were iso-C_16:0_ and anteiso-C_15:0_. The polar lipids were DPG, PE, PL (two) APL (three) and L (three).

*Paenibacillus* sp. S150: the strain was isolated from the wood of grapevine (*Vitis vinifera* L.), Sauvignon blanc cultivar in France. The genome size of this strain is 7.45 Mb with G+C content of 52.5 mol%. Genome sequencing and assembly BioProject is available on NCBI as PRJNA643435 (BioSample SAMN15414942). The assembly accession is available on GenBank and Refseq as GCA_019352175.1. The master record of the WGS project is JACBYI000000000. The GenBank accession of the 16S rRNA gene sequence is JACBYI010000087.

The strain was deposited in the collections of Institut Pasteur and DSMZ with the accession numbers XXXXXX and XXXXXX respectively.

## Acknowledgments

This work is supported by the Industrial Chair WinEsca funded by ANR (French National Research Agency) and the JAs Hennessy & Co and the GreenCell companies to PR. SC received funding via DaFNE Project Nr. 101384 from the Austrian Federal Ministry for Sustainability and Tourism (BMNT). We are grateful to B. Nikolic from AIT Austrian Institute of Technology for DNA isolation of strain S150.

## Conflicts of interest

The authors declare that there are no conflicts of interest.

## Notes

### Competing Interest Statement

The authors have declared no competing interest.

